# Prediction of Protein-Protein Interactions Based on L1-Regularized Logistic Regression and Gradient Tree Boosting

**DOI:** 10.1101/2020.03.04.976365

**Authors:** Bin Yu, Cheng Chen, Hongyan Zhou, Bingqiang Liu, Qin Ma

## Abstract

Protein-protein interactions (PPIs) are of great importance to understand genetic mechanisms, disease pathogenesis, and guide drug design. With the increase of PPIs sequence data and development of machine learning, the prediction and identification of PPIs have become a research hotspot in proteomics. In this paper, we propose a new prediction pipeline for PPIs based on gradient tree boosting (GTB). First, the initial feature vector is extracted by fusing pseudo amino acid composition (PseAAC), pseudo-position-specific scoring matrix (PsePSSM), reduced sequence and index-vectors (RSIV) and autocorrelation descriptor (AD). Second, to remove redundancy and noise, we employ L1-regularized logistic regression to select an optimal feature subset. Finally, GTB-PPI model based on GTB is constructed. Five-fold cross-validation showed GTB-PPI achieved the accuracies of 95.15% and 90.47% on *Saccharomyces cerevisiae* and *Helicobacter pylori*, respectively. In addition, GTB-PPI could be applied to predict *Caenorhabditis elegans*, *Escherichia coli*, *Homo sapiens,* and *Mus musculus* independent test sets, the one-core PPIs network for CD9, and the crossover PPIs network. The results show that GTB-PPI can significantly improve prediction accuracy of PPIs. The code and datasets of GTB-PPI can be downloaded from https://github.com/QUST-AIBBDRC/GTB-PPI/.

## Introduction

Knowledge of protein-protein interactions (PPIs) can help to grasp the mechanism of living cells, such as DNA replication, protein modification, and signal transduction [1,2]. The accurate understanding and analysis of PPIs can reveal multiple functions at the molecular and proteome level and has become a research hotspot [3,4]. However, web-lab identification methods suffer from incomplete, false prediction problem [5].

Therefore, using reliable bioinformatics methods can reduce research cost and play a guiding and complementary role. Compared with structure-based methods, the sequence-based methods are straightforward and does not involve prior information, which has been widely used. Martin et al. [6] proposed the signature kernel method to extract protein sequence feature information. The disadvantage was that Martin et al. did not use physicochemical property information. Guo et al. [7] employed seven physicochemical properties of amino acids to predict PPIs by combining auto covariance and support vector machine (SVM).

Different feature extraction methods can complement each other, and prediction accuracy can be improved by effective feature fusion [8,9]. Du et al. [8] constructed a PPI prediction framework called DeepPPI, which employed deep neural networks as the classifier. They fused amino acid composition information-based, physiochemical property-based sequence features. However, there may be redundant information and noise after feature fusion, and an excessively high dimension problem will affect the classification accuracy. You et al. [10] used the max-relevance-min-redundancy (mRMR) to determine important and distinguishable features to predict PPIs based on SVM.

Ensemble learning systems can achieve higher prediction performance than single classifier. To our knowledge, Jia et al. [11] combined seven random forest classifiers according to voting principles. Gradient tree boosting (GTB) is a good ensemble learning method and has been widely applied in bioinformatics, such as miRNA-disease association [12], drug-target interaction [13], RNA binding residue prediction [14]. The experimental results showed that GTB outperformed SVM, random forest (RF), and had a good model generalization performance.

Although a large number of computational technologies have been proposed and developed, there are still some challenges and problems in existing sequenced-based PPIs predictors. First, the sequence-only-based information of PPIs is not fully represented and elucidated, and satisfactory results cannot be obtained by adjusting individual parameters. Secondly, there is a severe data imbalance problem for the prediction of PPIs. The number of non-interaction pairs is much more significant than that of interaction protein pairs. At present, machine learning methods do not deal with such problems well. Statistical results show that machine learning methods result in poor overall performance when dealing with imbalanced data [15].

Inspired by the limits of machine learning methods, this paper proposes a new PPIs prediction pipeline called GTB-PPI based on GTB. First, we fuse pseudo amino acid composition (PseAAC), pseudo position-specific scoring matrix (PsePSSM), reduced sequence and index-vectors (RSIV) and autocorrelation descriptors (AD) to extract amino acid composition-based information, evolutionary information, and physicochemical information. To achieve effective details representing PPIs without losing important and reliable characteristic information, L1-regularized logistic regression is first utilized for PPIs prediction to eliminate redundant features. At the same time, we first employ GTB as a classifier to bridge the gap between the extracted PPI features and class label. Experimental results show that GTB outperforms the prediction performance of the SVM, RF, NB, and KNN classifiers. The linear combination of decision trees can fit the PPIs data well. When applied to the one-core for CD9 and crossover PPIs network, GTB-PPI obtains good generalization performance.

## Methods

### Data source

*S. cerevisiae* and *H. pylori* are applied to perform the pipeline of GTB-PPI. *S. cerevisiae* is from the DIP (the version of DIP_20070219) [7], where the amino acid sequence whose number of residues < 50 and sequence identity ≥ 40% via CD-HIT [16] are removed. Thus, 5,594 protein interaction pairs are positive samples, and 5,594 pairs with different subcellular location information are selected as negative samples.

Martin et al. [6] constructed *H. pylori*, which contains 2,916 samples (1,458 PPIs pairs and 1,458 non-PPIs pairs). Four independent PPIs datasets [17] are also used to verify the GTB-PPI, including *Caenorhabditis elegans* (*C. elegans*, 4,013 interacting pairs), *Escherichia coli* (*E. coli*, 6,954 interacting pairs), *Homo sapiens* (*H. sapiens*, 1,412 interacting pairs), and *Mus musculus* (*M. musculus*, 313 interacting pairs). The number of unique proteins in each dataset are shown in Table S1.

### Feature extraction

In the GTB-PPI pipline, we fuse PseAAC, PsePSSM, RSIV, AD to obtain the PPIs feature information. PseAAC has been applied in PPIs prediction [18], Gram-negative protein localization prediction [19] and identification of submitochondrial locations [20]. And PsePSSM has been employed to locate the apoptosis proteins [21]. Four feature extraction descriptors can elaborate amino acid composition-based features, physicochemical property features, and evolutionary information features associated with PPIs. The detained descriptions of methods are in File S1.

### L1-regularization logistic regression

The L1-regularized logistic regression (L1-RLR) is an embedded feature selection method. Given the sample data set *D* = {(*x*_1_, *y*_1_),(*x*_2_, *y*_2_),…,(*x*_*m*_, *y*_*m*_)}. Thus, the L1-RLR can be transformed into an unconstrained optimization problem.

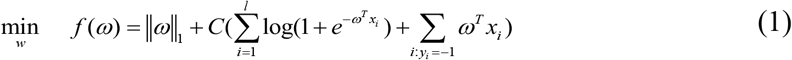

where ||·||_1_ represents the L1 norm. *ω* represents the weight coefficient. *C* represents penalty term, which determines the number of selected features. We use the coordinate descent algorithm in LIBLINEAR [22] to solve Equation (1).

### Gradient tree boosting

Gradient tree boosting (GTB) is an algorithm that aggregates multiple decision trees [23,24]. The difference between GTB and other ensemble learning algorithms is that GTB fits residual of the regression tree at each iteration using negative gradient values of loss.

GTB can be expressed as the relationship between the label *y* and the vector of input variables *x* connected via a joint probability distribution *p*(*x*, *y*). The goal of GTB is to obtain the estimated function 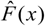 through minimizing *L*(*y*, *F*(*x*)):

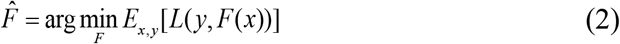

Let *h*_*m*_(*x*) be the *m*-*th* decision tree. *J*_*m*_ indicate number of its leaves. The tree partitions the input space into *J*_*m*_ disjoint regions 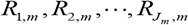 and predicts a numerical value *b*_*jm*_ for each region *R*_*jm*_. The output of *h*_*m*_(*x*) can be described as:

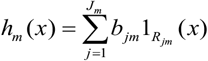

Then we can obtain *γ*_*m*_ value using steepest descent to fulfill the GTB model:

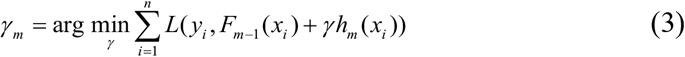

where *F*_*m*−1_(*x*) represents an estimated function. The iterative criterion of GTB is shown using Equation (4).

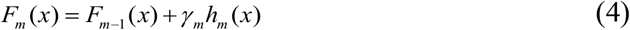

Iterations are set as *M*, and gradient tree boosting model is 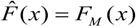.

The GTB can complement the weak learning ability of decision tree, improving the ability of representation, optimization, and generalization, which can capture higher-order information and is invariant to scaling of sample data. GTB can effectively avoid overfitting condition by weighting combination scheme. GTB-PPI uses the GTB algorithm of Scikit-learn [25].

### Performance evaluation

In GTB-PPI pipline, Recall, Precision, Overall Prediction Accuracy (ACC), and Matthews correlation coefficient (MCC) is taken into the count to evaluate the model performance [8]. The definitions are as follows:

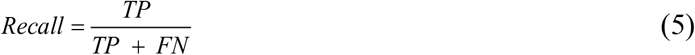

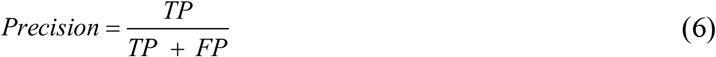

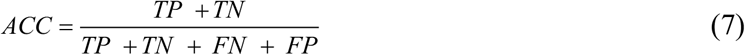

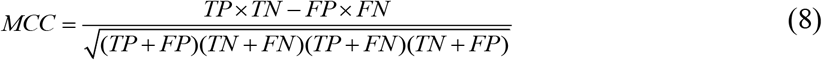

TP indicates the number of predicted PPI samples found in PPI dataset; TN indicates the number of non-PPIs samples correctly predicted; FP, FN indicates false positive, and false negative. Receiver Operating Characteristic (ROC) curve [26], Precision-Recall (PR) curve [27], the area under ROC curve (AUC) and PR curve (AUPR) are also used to evaluate the generalization ability of the GTB-PPI.

## Results

This paper proposes a PPIs prediction method based on GTB called GTB-PPI. The pipeline of GTB-PPI for predicting PPIs is shown in **Figure 1**.

**Figure 1.**
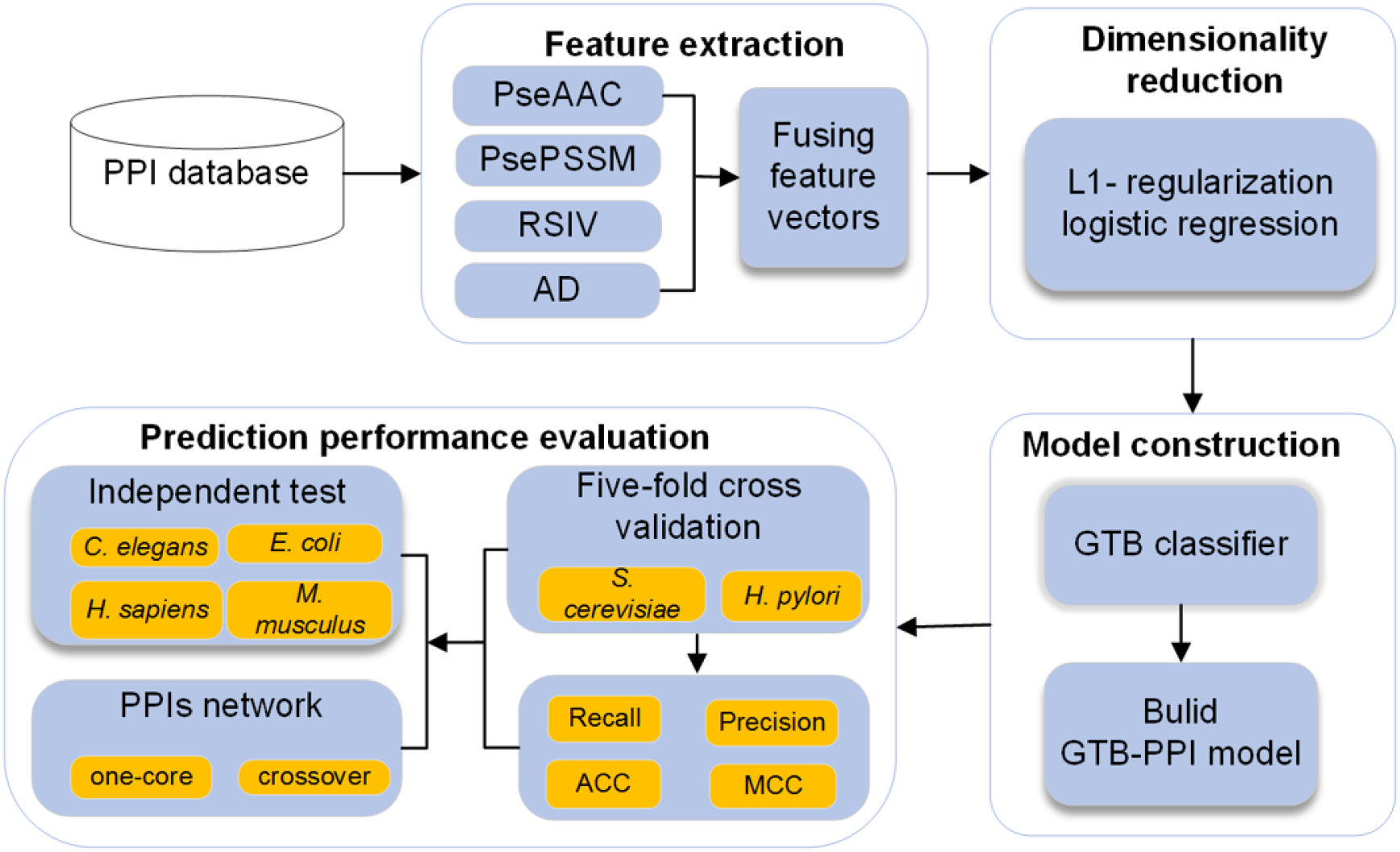
Overall framework of GTB-PPI for protein-protein interactions prediction. First, PseAAC, PsePSSM, RSIV and AD are used for feature extraction and the L1-regularization logistic regression is employed for dimensionality reduction. Secondly, we use GTB to predict PPIs and GTB-PPI model is constructed. Finally, five-fold cross validation and independent test are employed to evaluate GTB-PPI.

The steps of GTB-PPI can be described as:

1) Input the PPI samples, non-PPIs samples and the corresponding binary labels.

2) Feature extraction. Fusing PseAAC, PsePSSM, RSIV, and AD to transform the protein character signal into numerical signal. (a) Obtain amino acid sequence composition and sequence order information using the PseAAC. Thus, the 20 + *λ* dimensional vectors are constructed. (b) Obtain the PSSM matrix of the protein sequence and extract 20 + 20×*ξ* features based on PsePSSM. (c) According to the six physicochemical properties, the feature information is extracted using RSIV. Each protein sequence constructed a 120+77=197 dimensional vector. (d) The protein sequence is transformed into a 3× 7 × *lag* dimensional vector by NMBA, MA, and GA.

3) Dimensionality reduction. (a) The L1-RLR is employed to remove redundant features by adjusting the penalty parameters in logistic regression. (b) The L1-RLR is compared with semi-supervised dimension reduction, principal component analysis, kernel principal component analysis, factor analysis, minimal-redundancy-maximal-relevance, and conditional mutual information maximization on *S. cerevisiae* and *H. pylori*.

4) Predict PPIs based on GTB. According to step 2) and step 3), using L1-RLR can better capture the sequence representation details. In this way, GTB-PPI model can be constructed using GTB as the classifier.

5) Predict PPIs on independent test sets and network data. Feature encoding, fusion, and selection can effectively provide the PPIs representation. Then GTB is employed to predict the binary labels on four independent test sets and two network sets.

### Parameter optimization of PseAAC, PsePSSM, and AD

The parameter optimization of PseAAC, PsePSSM, and AD plays an essential role in GTB-PPI predictor construction. We implement the hyperparameter optimization through five-fold cross-validation. The pipeline of GTB-PPI is implemented using MATLAB 2014a and Python 3.6.

When we extract features from the sequence, the *λ* of PseAAC, the *ξ* of PsePSSM, and the *lag* of AD should be determined. We set the *λ* values as 1, 3, 5, 7, 9, and 11 and the *ξ* and *lag* are also set as 1, 3, 5, 7, 9, and 11 in order. The GTB is used to predict the binary label (Table S4, Table S5, and Table S6). We also plot **Figure 2** to show the effect of selecting different values of parameters intuitively.

**Figure 2.**
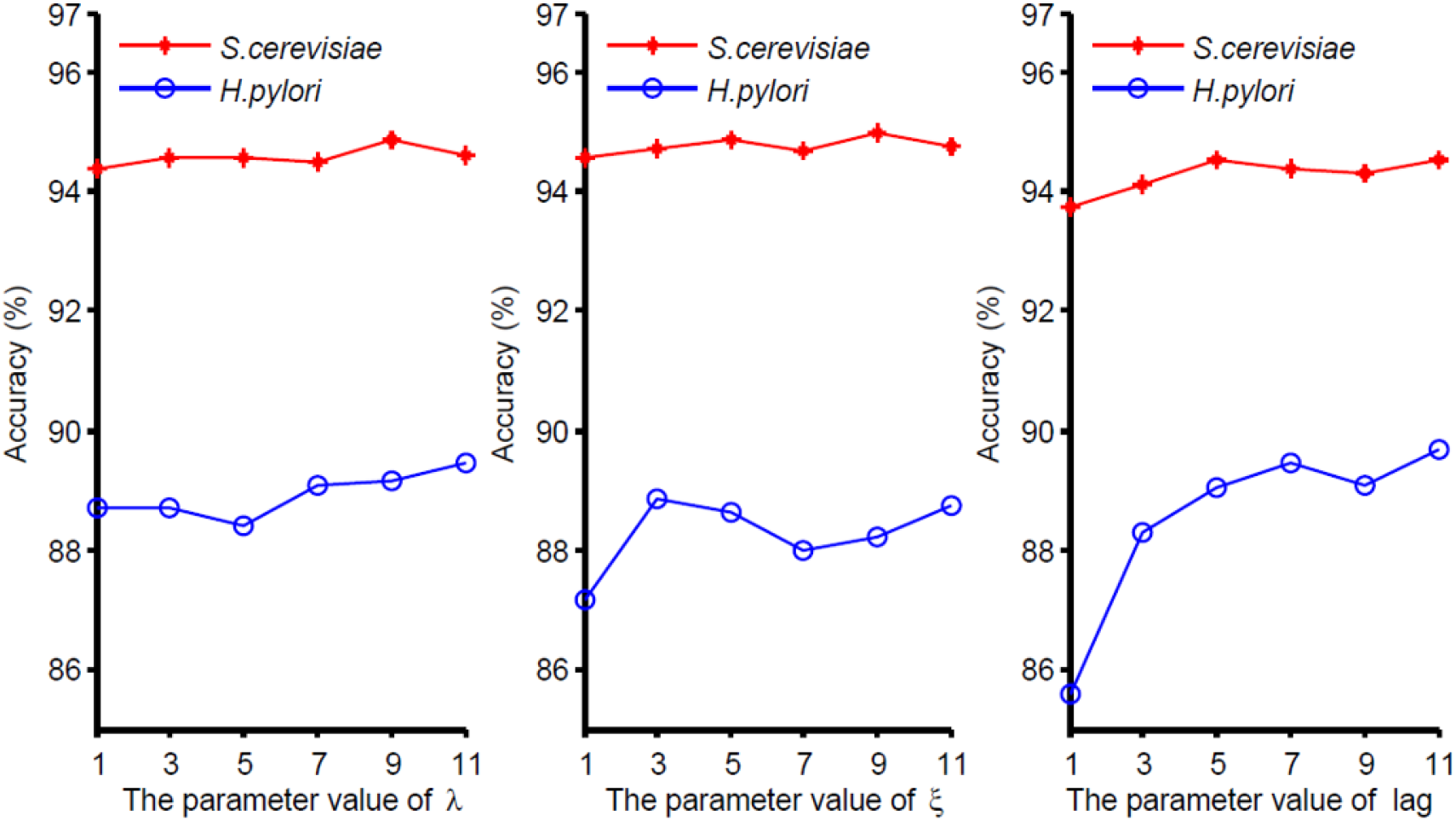
Prediction results of different parameters *λ*, *ξ* and *lag* on *S. cerevisiae* and *H. pylori* datasets. The *λ*, *ξ* and *lag* are the parameters need to be adjusted in PseAAC, PsePSSM and AD.

From **Figure 2**, we can see that with the change of the parameter’s value, the prediction performance of *S. cerevisiae* and *H. pylori* are also changing. For the parameter *λ* in PseAAC, *λ* values are different when two datasets achieve the highest prediction performance. The optimal *λ* value for *S. cerevisiae* is 9, while the optimal *λ* value of *H. pylori* is 11. Considering that PseAAC generates less dimensional vectors than the other three feature extraction methods, we choose the optimal parameter *λ*=11 to mine more PseAAC information. The discussion of *ξ* and *lag* can be find in File S2. In conclusion, for each protein sequence, PseAAC extracts 20 +11 = 31 features, PsePSSM obtains 20+20×9=200 features, the dimension of RSIV is 197, and AD encodes 3×7×11 = 231 features. We can obtain 659-dimensional vector by fusing four coding methods. Then the 1318-dimensional feature vector is constructed by concatenating two sequences of protein pairs.

### The effect of dimensionality reduction

The L1-RLR can effectively improve predictive performance with higher computational efficiency. The process of papameter selection is set in File S3. In order to better evaluate the merits of L1-RLR (*C* = 1), we compared it with semi-supervised dimensionality reduction (SSDR) [28], principal component analysis (PCA) [29] (The setting of contribution rate is shown in Table S8), kernel principal component analysis (KPCA) [30] (The adjustment of contribution rate is shown in Table S9), factorial analysis (FA) [31], mRMR [32], and conditional mutual information maximization (CMIM) [33] (Table S10). The ROC and PR curves of different dimensional reduction methods are plotted, which are shown in **Figure 3**. The AUC and AUPR are shown in Table S11.

**Figure 3.**
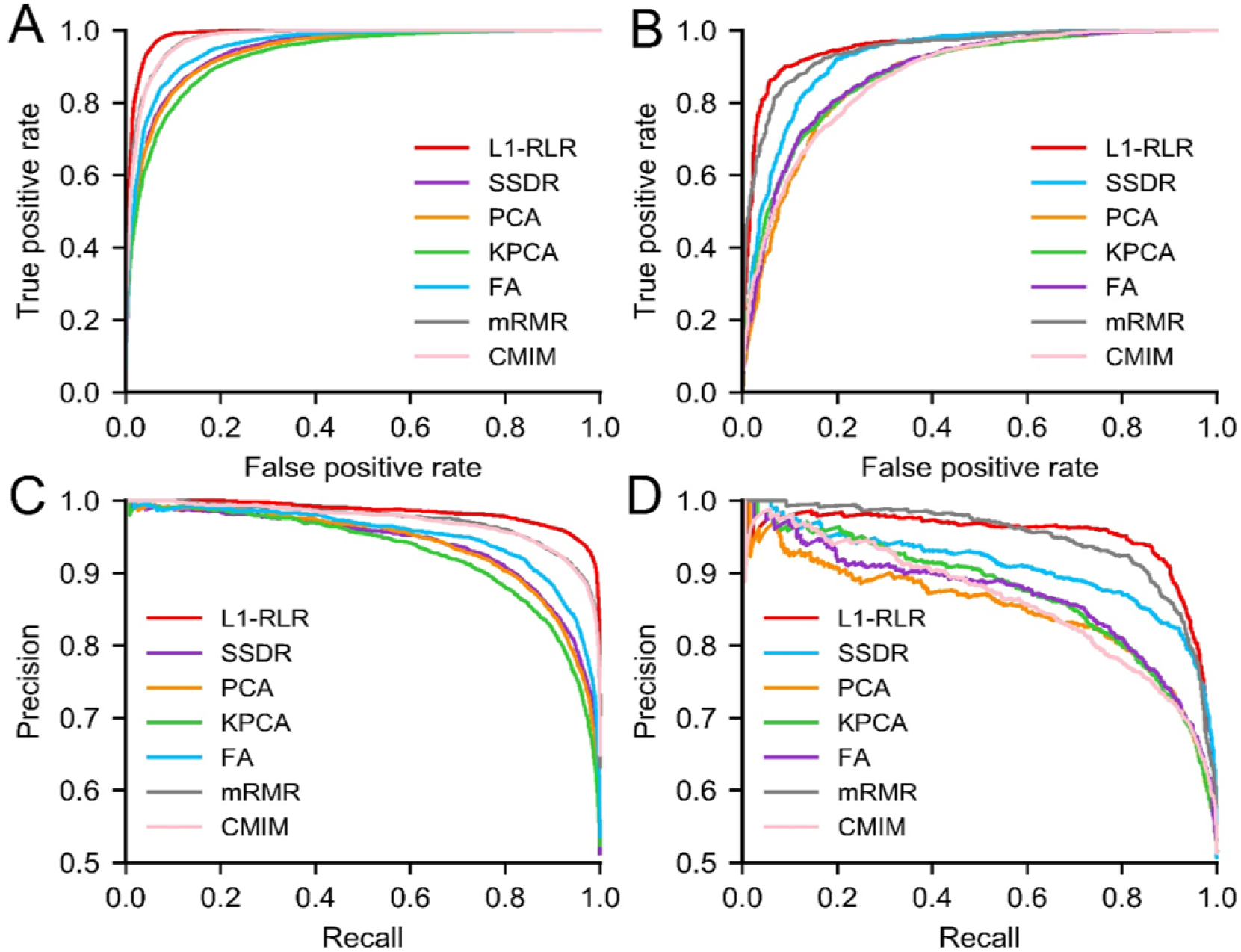
Predictive performance of different dimensional reduction methods. **A-B.** The ROC curves indicate L1-RLR achieves the higher AUC than SSDR, PCA, KPCA, FA, mRMR, and CMIM on *S. cerevisiae* and *H. pylori*. **C-D.** The PR curves of *S. cerevisiae* and *H. pylori* show L1-RLR obtains the superior AUPR.

From Figure 3A, ROC curves show that the L1-regularization logistic regression has superior model performance. For *S. cerevisiae*, the AUC values of L1-RLR, SSDR, PCA, KPCA, FA, mRMR, and CMIM are 0.9875, 0.9445, 0.9420, 0.9305, 0.9568, 0.9770, and 0.9769, respectively. The AUC value of L1-RLR is 4.3% higher than SSDR. Figure 3B plots the ROC curves on the *H. pylori* illustrating the comparison of different dimensional reduction methods. We can know the AUC values of L1-RLR, SSDR, PCA, KPCA, FA, mRMR, and CMIM are 0.9559, 0.9238, 0.8706, 0.8803, 0.8824, 0.9461, and 0.8726, respectively. The AUC value of L1-RLR is 3.21%, 8.53%, 7.56%, 7.35%, 0.98%, and 8.33% higher than SSDR, PCA, KPCA, FA, mRMR, and CMIM respectively. The analysis of PR curves are shown in File S4. In the field of PPIs, the L1-regularized logistic regression is first used to reduce the dimension, which could remove the redundant features without hurting important information. The effective features related to PPIs could be fed into a GTB classifier, generating a reliable GTB-PPI prediction model.

### The selection of classifier algorithms

GTB is used as a classifier whose number of iterations is set to 1000, and the loss function is set as “deviance”. The main results of other classifiers are also provided via five-fold cross-validation, including *K* nearest neighbors (KNN) [34] whose number of neighbors is three (Table S12), Naïve Bayes (NB) [35], SVM [36] whose kernel function is RFE, and RF [37] whose number of the base decision tree is 1000 (Table S13), which are shown in Table S14. The histograms of KNN, SVM, NB, RF, and GTB on *S. cerevisiae* and *H. pylori* are shown in Figure S3 and Figure S4, respectively. We also obtain the ROC and PR curves (**Figure 4**), and AUC value and AUPR value of different classifiers (Table S15).

**Figure 4.**
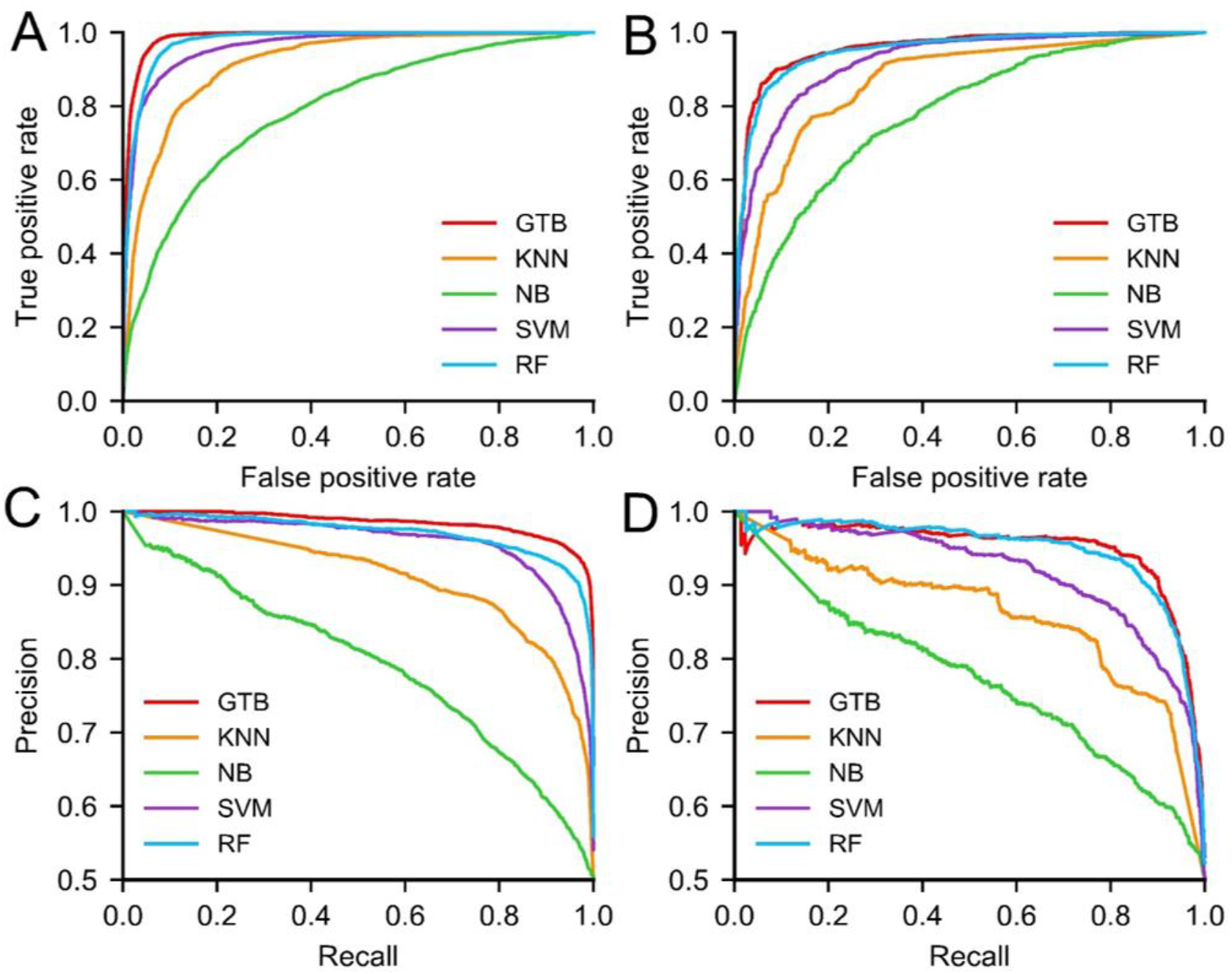
Comparison of GTB with KNN, NB, SVM, and RF classifiers. **A-B.** The ROC curves of *S. cerevisiae* and *H. pylori* indicate GTB achieves superior AUC value than KNN, NB, SVM, and RF. **C-D.** The PR curves illustrating prediction performance of different classifiers on *S. cerevisiae* and *H. pylori*.

The comparison of different classifiers (File S5) shows that the GTB outperforms prediction performance of SVM, RF, NB, and KNN classifiers. GTB-PPI can accurately indicate whether a protein pair interact on *S. cerevisiae* and *H. polyri*. The GTB complements the weakness of the decision tree’s learning ability, avoiding overfitting and showing excellent performance. RF is an ensemble classifier using Bagging algorithm. But all the base decision trees are treated equally. If the base classifier’s prediction performance is biased, the final ensemble classifier may get the unreliable and biased predicted results. GTB is an ensemble learning algorithm that can achieve superior generalization performance over single learner, which can effectively bridge the gap betwwen the sequence and PPIs label information via steepest descent step algorithm.

## Discussion

### Comparison with other PPIs prediction methods

To verify the validity of the GTB-PPI model, We compare GTB-PPI with ACC+SVM [7], DeepPPI [8], and other state-of-the-art methods on *S. cerevisiae* and *H. pylori*, which are shown in **Table 1** and **Table 2**. In order to further evaluate the performance on cross-species datasets based on GTB-PPI model. The *S. cerevisiae* is used as training set, and four independent test sets to predict PPIs. The main results are shown in **Table 3**.

**Table 1.**
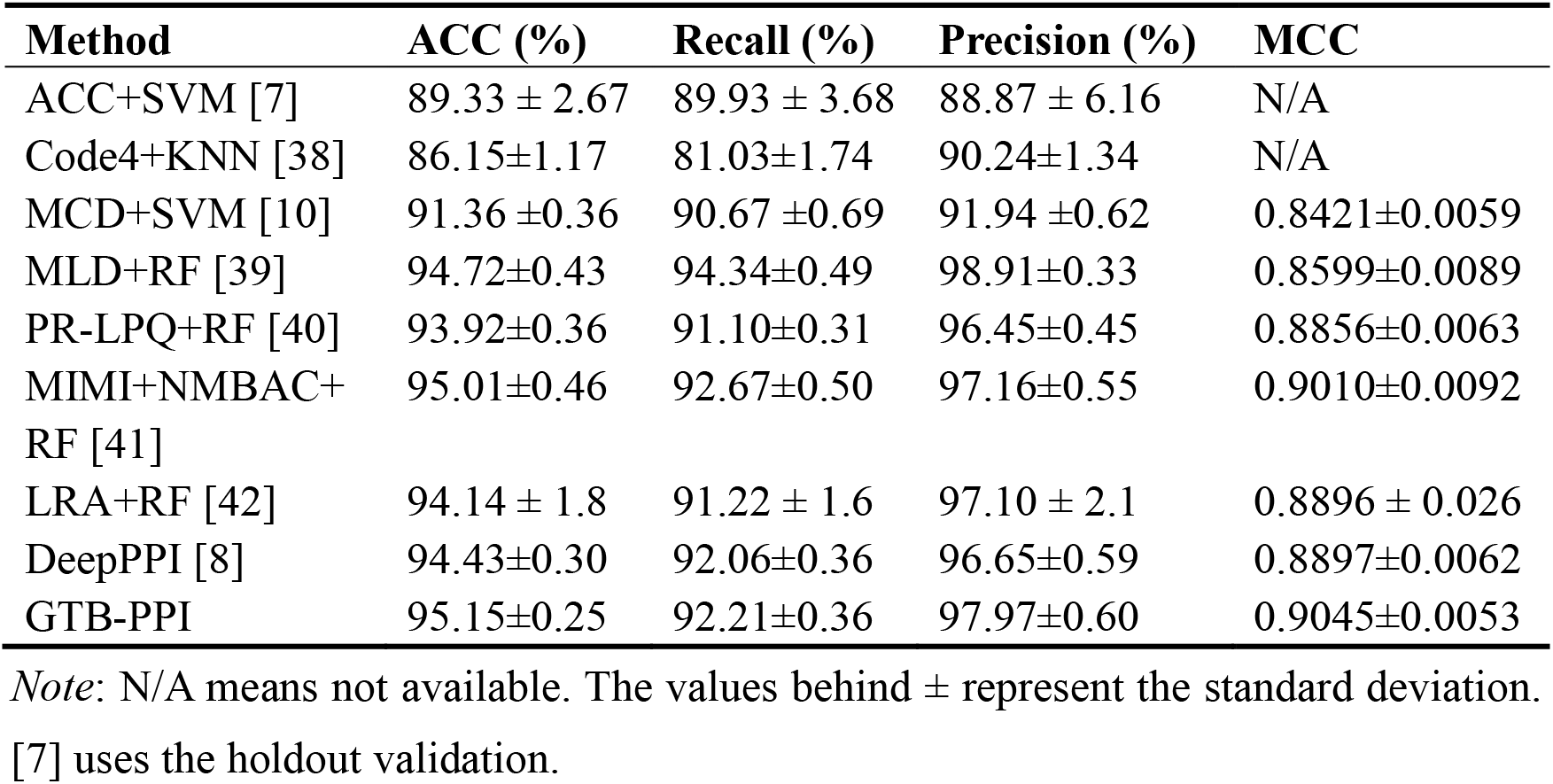
Comparison with other state-of-the-art predictors on *S. cerevisiae*.

| Method | ACC (%) | Recall (%) | Precision (%) | MCC |
| --- | --- | --- | --- | --- |
| ACC+SVM [7] | $89.33 \pm 2.67$ | $89.93 \pm 3.68$ | $88.87 \pm 6.16$ | N/A |
| Code4+KNN [38] | $86.15 \pm 1.17$ | $81.03 \pm 1.74$ | $90.24 \pm 1.34$ | N/A |
| MCD+SVM [10] | $91.36 \pm 0.36$ | $90.67 \pm 0.69$ | $91.94 \pm 0.62$ | $0.8421 \pm 0.0059$ |
| MLD+RF [39] | $94.72 \pm 0.43$ | $94.34 \pm 0.49$ | $98.91 \pm 0.33$ | $0.8599 \pm 0.0089$ |
| PR-LPQ+RF [40] | $93.92 \pm 0.36$ | $91.10 \pm 0.31$ | $96.45 \pm 0.45$ | $0.8856 \pm 0.0063$ |
| MIMI+NMBAC+RF [41] | $95.01 \pm 0.46$ | $92.67 \pm 0.50$ | $97.16 \pm 0.55$ | $0.9010 \pm 0.0092$ |
| LRA+RF [42] | $94.14 \pm 1.8$ | $91.22 \pm 1.6$ | $97.10 \pm 2.1$ | $0.8896 \pm 0.026$ |
| DeepPPI [8] | $94.43 \pm 0.30$ | $92.06 \pm 0.36$ | $96.65 \pm 0.59$ | $0.8897 \pm 0.0062$ |
| GTB-PPI | $95.15 \pm 0.25$ | $92.21 \pm 0.36$ | $97.97 \pm 0.60$ | $0.9045 \pm 0.0053$ |

Note: N/A means not available. The values behind ± represent the standard deviation. [7] uses the holdout validation.

**Table 2.** Comparison with other state-of-the-art predictors on *H. pylori*.

| Method | ACC (%) | Recall (%) | Precision (%) | MCC |
| --- | --- | --- | --- | --- |
| SVM [6] | 83.40 | 79.90 | 85.70 | N/A |
| WSR [43] | 83.70 | 79.00 | 87.00 | N/A |
| Ensemble of HKNN [44] | 86.60 | 86.70 | 85.00 | N/A |
| DCT+WSRC [45] | 86.74 | 86.43 | 87.01 | 0.7699 |
| MCD+SVM [10] | 84.91 | 83.24 | 86.12 | 0.7440 |
| MIMI+NMBAC+RF [41] | 87.59 | 86.81 | 88.23 | 0.7524 |
| DeepPPI [8] | 86.23 | 89.44 | 84.32 | 0.7263 |
| GTB-PPI | 90.47±0.84 | 91.15±1.42 | 89.99±2.06 | 0.8100±0.0163 |

Note: N/A means not available. The values behind ± represent the standard deviation. [6], [43] and [44] use ten-fold cross-validation.

**Table 3.** Comparison of performance of the proposed method with other state-of-the-art predictors on the independent dataset.

| Species | ACC (%) |  |  |  |  |
| --- | --- | --- | --- | --- | --- |
|  | GTB-PPI | DeepPPI [8] | MIMI+NMB AC+RF [41] | MLD+RF [39] | DCT+WSR C [45] |
| *C. elegans* | 92.42 | 94.84 | 92.16 | 87.71 | 81.19 |
| *E. coli* | 94.06 | 92.19 | 92.80 | 89.30 | 66.08 |
| *H. sapiens* | 97.38 | 93.77 | 94.33 | 94.19 | 82.22 |
| *M. musculus* | 98.08 | 91.37 | 95.85 | 91.96 | 79.87 |

As shown in **Table 1**, compared with other existing methods listed, the ACC of GTB-PPI increased by 0.1%-9%. The ACC of GTB-PPI is 0.72% higher than Du et al. [8], 1.23% higher than the Wong et al. [40] prediction method, and 1.01% higher than You et al. [42]. The Recall of GTB-PPI is 0.15% higher than Du et al. [8] and 1.54% higher than You et al. [10]. The Precision of GTB-PPI is 1.32% higher than Du et al. [8] and 0.81% higher than Ding et al. [41].

As shown in **Table 2**, the performance of GTB-PPI is better than the most accurate predictors. The ACC, Recall, Precision, and MCC values obtained are 90.47%, 91.15%, 89.99% and 0.8100, respectively. In terms of ACC, GTB-PPI is 2.88%-7.07% higher than other methods, 7.07% higher than Martin et al., 4.24% higher than Du et al. [8] and 3.73% higher than Huang et al. [45]. At the same time, the Recall of GTB-PPI is 1.71%-12.15% higher than other methods, 4.72% higher than Huang et al. [45] and 7.91% higher than You et al. [10]. The Precision of GTB-PPI is 1.76%-5.67% higher than other methods, 4.29% higher than Martin et al. [6], and 5.67% higher than Du et al. [8].

From **Table 3**, for the *C. elegans*, GTB-PPI is 0.26% higher than Ding et al. [41], 4.71% higher than You et al. [39], and 11.23% higher than Huang et al. [45]. For the *E. coli*, the ACC of GTB-PPI is 1.26%-27.98% higher than DeepPPI (92.19%) [8], MIMI+NMBAC+RF (92.80%) [41], MLD+RF (89.30%) [39], DCT+WSRC (66.08%) [45]. For the *H. sapiens*, GTB-PPI is 3.05%-15.16% higher than DeepPPI (93.77%) [8], MIMI+NMBAC+RF (94.33%) [41], MLD+RF (94.19%) [39], DCT+WSRC (82.22%) [45]. For the *M. musculus*, GTB-PPI is 2.23%-18.21% higher than the other PPIs predictors. The findings indicate the biological hypothesis of mapping PPIs from one species to another species is reasonable. Because PPIs in one organism might have “co-evolve” with other organisms [41].

### PPIs network prediction

The graph visualization of the PPIs network can provide broad and informative idea to understand the proteome and analyze the protein function. We employ GTB-PPI to predict the simple one-core PPIs network for CD9 [46] and PPIs crossover network for the Wnt-related pathway [47] using *S. cerevisiae* dataset as training set (**Figure 5**). The blue lines represent true prediction of PPIs, and the red lines represent false prediction of PPIs.

**Figure 5.**
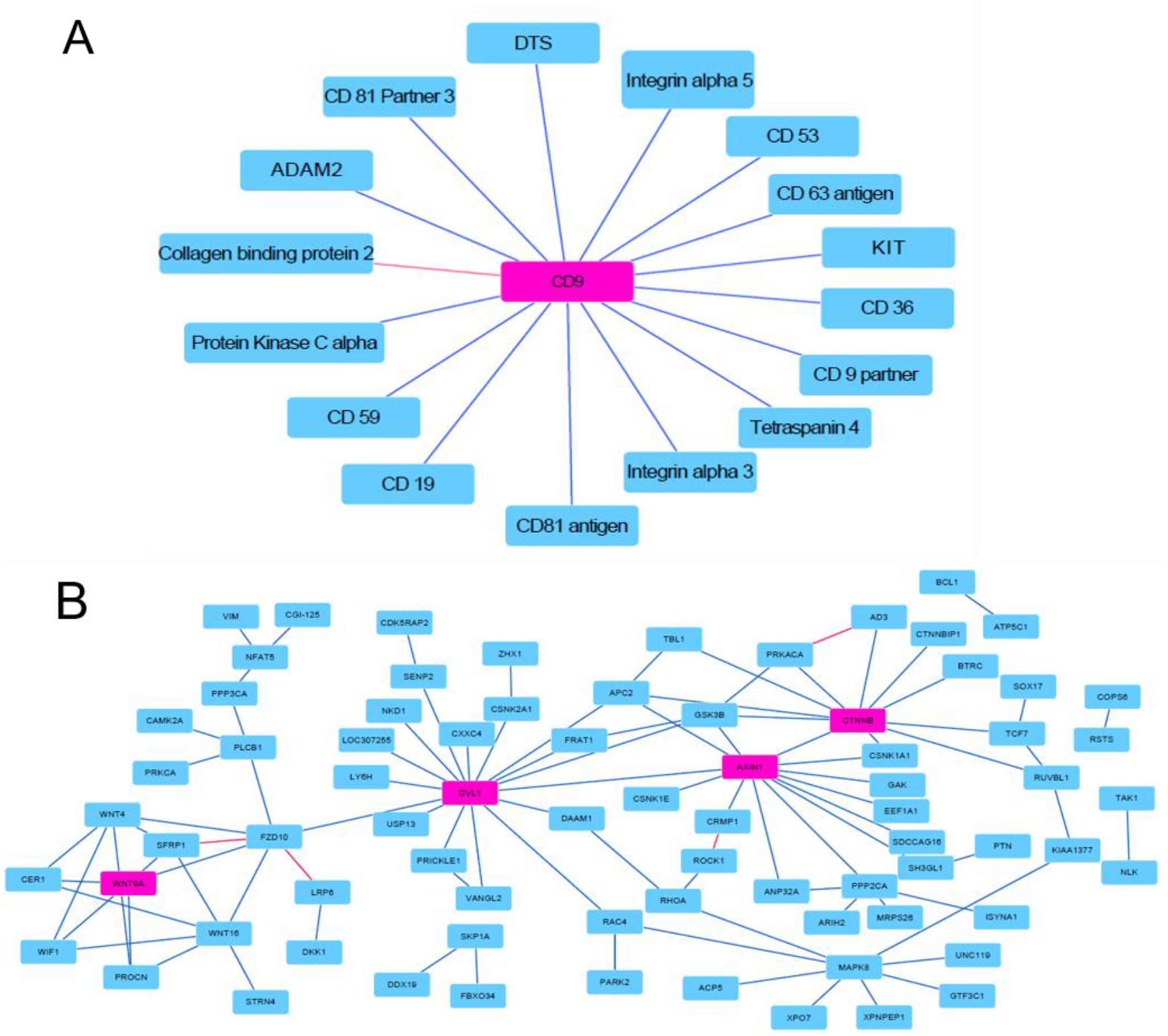
Predicted results of one-core network and crossover network. **A**. The prediction performance of one-core network. The CD9 is the core protein and 15 of all 16 PPIs are predicted successfully. **B**. The prediction performance of crossover network. The important proteins WNT9A, DVL1, AXIN1 and CTNNB are linked to the Wnt-related pathway. 92 interactions among the 96 PPIs pairs are identified based on GTB-PPI model.

As shown in Figure 5A, we can see only the interaction between CD9 and Collagen-binding protein 2 is not predicted successfully, and Shen et al. also cannot predict the interaction. Comparing with Shen et al. [48] and Ding et al. [41], GTB-PPI achieves the superior prediction performance. The overall accuracy is 93.75%, which is 12.5% higher than Shen et al. (81.25%) [48] and 6.25% higher than Ding et al. (87.50%) [41]. Figure 5B reveals that the overall accuracy of our method is 95.83%, which is 19.79% higher than Shen et al. (76.04%) [48] and 1.04% higher than Ding et al. (94.79%) [41].

The palmitoylation of CD9 could support CD9 associate with CD81 and CD 53 [49]. In the one-core network for CD9, we can see the interaction between CD 9 and CD 53 is predicted successfully based on GTB-PPI. In the crossover PPIs network for the Wnt-related signaling pathway, the ANP32A, CRMP1, and KIAA1377 are linked to the Wnt pathway via proteins. The ANP32A has been demonstrated as a potential tumor suppressor [50], and GTB-PPI could predict the interactions between the corresponding proteins. However, the interaction between ROCK1 and CRMP1 is not predicted. It is likely because we use *S. cerevisiae* as training set, and ROCK1, CRMP1 are different species from *S. cerevisiae*. At the same time, ROCK1 is part of the noncanonical Wnt signaling pathway [47], GTB-PPI may not be very effective in this case. AXIN1 could interact with multiple proteins [51]. GTB-PPI can predict the interactions between AXIN1 and its satellite proteins, which provides a new thought to elucidate the biological function and to expedite PPIs network.

## Conclusion

The knowledge and analysis of PPIs can help us to reveal the structure and function at the molecular level, such as growth, development, metabolism, signal transduction, differentiation, and apoptosis. In this paper, a new protein-protein interaction prediction pipline called GTB-PPI is presented. First, PseAAC, PsePSSM, RSIV, and AD are concatenated as the initial feature information for predicting PPIs. PseAAC obtains not only the amino acid composition information but also the order sequence information. PsePSSM can mine the evolutionary information and local order information, while RSIV could obtain the frequency feature information using the reduced sequence. AD reflects the physicochemical property feature on global amino acid sequence. Secondly, L1-regularized logistic regression could obtain effective information features related to PPIs without losing accuracy and generalization. And L1-RLR is superior to SSDR, PCA, KPCA, FA, mRMR, and CMIMs. Finally, the PPIs are predicted based on GTB which employs decision tree as the base classifier, which can bridge the gap betwwen amino acid sequence information features and class label. Experimental results show that the GTB outperforms the SVM, RF, NB, and KNN classifiers. Especially, in the field of binary PPIs prediction, the L1-regularized logistic regression is used for dimensionality reduction for the first time. The gradient tree boosting is also first employed as a classifier. In a word, GTB-PPI shows good performance, representation ability, and generalization ability.

## Data policy

The all datasets and code of GTB-PPI can be obtained on https://github.com/QUST-AIBBDRC/GTB-PPI/. The Dataset file contains eight PPIs datasets.

## Authors’ contributions

BY, CC, and QM conceived the algorithm, prepared the datasets, carried out experiments, and wrote the manuscript. HZ and BL designed, performed and analyzed experiments. All authors revised the manuscript. All authors read and approved the final manuscript.

## Competing interests

The authors have declared that they have no competing interests.

## Acknowledgments

This work was supported by the National Natural Science Foundation of China (No. 61863010), the Key Research and Development Program of Shandong Province of China (2019GGX101001), and the Natural Science Foundation of Shandong Province of China (Nos. ZR2017MA014 and ZR2018MC007). This work used the Extreme Science and Engineering Discovery Environment, which is supported by the National Science Foundation (No. ACI-1548562).

## Supplementary material

**File S1 Feature extraction methods**

**File S2 The parameter selection of *ξ* and *lag***

**File S3 The parameter optimization of L1-RLR**

**File S4 The comparison of dimensional-reduction methods**

**File S5 The comparison of different classifiers**

## Supplementary figure legends

**Figure S1 The number of feature subset with different feature extraction methods on *S. cerevisiae* dataset**

The raw features represent the number of initial feature dimension and the optimal features represent the number of the selected features.

**Figure S2 The number of feature subset with different feature extraction methods on *H. pylori* dataset**

The raw features represent the number of initial feature dimension and the optimal features represent the number of the selected features.

**Figure S3 The comparison of prediction results with different classifiers on *S. cerevisiae* dataset**

**Figure S4 The comparison of prediction results with different classifiers on *H. pylori* dataset**

## Supplementary tables

**Table S1 The unique proteins for each dataset**

**Table S2 Amino acid classification according to different physicochemical property**

**Table S3 The original values of the seven physicochemical properties for the 20 native amino acids**

**Table S4 Performance comparison with different *λ* values on PPIs datasets**

**Table S5 Performance comparison with different *ξ* values on PPIs datasets**

**Table S6 Prediction results with the different *lag* value on PPIs data set**

**Table S7 Effect of selecting different penalty parameter on the model performance**

**Table S8 Performance of kernel principle component analysis with different contribution rate**

**Table S9 Performance of principle component analysis with different contribution rate**

**Table S10 Comparison of prediction results on different dimensional reduction methods**

**Table S11 The AUC and AUPR on different dimensional reduction methods**

**Table S12 Performance of K nearest neighbor with different size of neighbor**

**Table S13 Performance of random forest with different size of base decision tree**

**Table S14 Prediction results of different classifiers on *S. cerevisiae*, *H. pylori* dataset**

**Table S15 The AUC and AUPR of different classifiers on *S. cerevisiae*, *H. pylori* dataset**

