## Supplementary files, Supplementary figures, Supplementary tables for "Prediction of Protein-Protein Interactions Based on L1-Regularized Logistic Regression and Gradient Tree Boosting"

#### File S1 Feature extraction methods

*Pseudo amino acid composition*

To extract the frequency and position information, Chou et al. [1] proposed a PseAAC method, which not only considered the composition of the protein sequence but also represented the positional sequence information.

The feature vector of PseAAC model can be expressed as:

$$X = [x_1, x_2, \cdots, x_{19}, x_{20}, x_{20+1}, \cdots, x_{20+\lambda}]^T \ (\lambda < L) \tag{1}$$

where $L$ indicates the sequence length.

$$x_u = \begin{cases} \dfrac{f_u}{\sum_{i=1}^{20} f_i + \omega \sum_{j=1}^{\lambda} \theta_j}, (1 \le u \le 20) \\ \dfrac{\omega \theta_{u-20}}{\sum_{i=1}^{20} f_i + \omega \sum_{j=1}^{\lambda} \theta_j}, (20+1 \le u \le 20+\lambda) \end{cases} \tag{2}$$

where $f_u$ is the normalized frequency of *u-th* amino acid and $\theta_j$ is the *j-th* sequence correlation factor. $\omega$ is set to 0.05 [1]. Equation (2) showed the dimension of the feature vector extracted by PseAAC is $20+\lambda$. The first 20 dimensions of the feature vector represent the amino acid composition information, whereas the components of the remaining $\lambda$ dimension reflect the information of sequence order. Since the shortest length of the sequence in the protein-protein interaction dataset is 12, the parameter must satisfy $\lambda < 12$.

*Pseudo position-specific scoring matrix*

Evolutionary information embedded in the PSSM has been applied in subcellular location prediction [2]. We used the PSI-BLAST tool [3] to search the non-redundant database Swiss-Prot to obtain PSSM of the protein sequence. During the running process, the PSI-BLAST parameter e-value threshold and the maximum number of iterations is set as 0.001 and 3, respectively.

First, the elements of the PSSM matrix are transformed into the [0, 1] interval via $f(x)$ sigmoid function.

$$f(x) = \frac{1}{1+e^{-x}} \tag{3}$$

To make PSSM become a size-uniform vector. The protein sample can be expressed

$$\bar{P}_{PSSM} = \bar{p}_1 \bar{p}_2 \cdots \bar{p}_{20} \tag{4}$$

where $\bar{p}_j = \frac{1}{L}\sum_{i=1}^{L} p_{i,j}$ represents the average score of the $j$ amino acid. However, only using $\bar{P}_{PSSM}$ could discard order information of PSSM. According to the pseudo amino acid composition, which was proposed by Shen and Chou [4], we obtain the PsePSSM feature vector by Equation (5).

$$\bar{p}_j = \begin{cases} \frac{1}{L}\sum_{i=1}^{L} a_{i,j} & j = 1, 2, \cdots, 20,\ \zeta = 0 \\ \frac{1}{L-\zeta}\sum_{i=1}^{L-\zeta} (a_{i,j} - a_{i+\zeta,j})^2 & j = 1, 2, \cdots, 20,\ \zeta < L \end{cases} \tag{5}$$

From Equation (5), each protein sequence can generate $20{+}20{\times}\xi$ the dimensional feature vector. The first 20-dimensional vector represents the composition information of the PSSM, and the remaining $20{\times}\xi$ dimensional feature vector represents the order of evolutionary information. PsePSSM can transform an inconsistent protein sequence into a consistent numerical vector.

*Reduced sequence and index-vectors*

The reduced alphabet schemes can effectively provide valuable information to represent PPIs. Xu et al. [5] extracted the amino acid sequence feature information using reduced sequence and index-vectors (RSIV) combined with q-FP and CMV feature extraction methods to predict therapeutic peptides. The amino acid sequence group coding in RSIV is shown in Table S2.

For instance, 20 amino acid residues are classified into four groups according to DHP. A represents {P, A, L, V, I, F, W, M}; B represents {Q, S, T, Y, C, N, G}, C represents {H, K, R}, D represents {D, E}.

For protein {MPNDNKTPNRSSTPKFTKKPVTPNDKIPEREEKSN}. The reduced sequence is {AABDBCBABCBBBACABCCAABABDCAADCDDCBB}. In the reduced sequence, the frequency of 'A' is 10; the 'B' is 12, the 'C' is 8, and the 'D' is 5. The frequency of di-character 'AA' is 3; the frequency 'AB' is 5; ⋯; the frequency 'AD' is 1. And in this case, the frequency of 'DD' is 1.

Thus, RSIV can generate two types of index-vectors $V_1$ and $V_2$, which are shown in Equation (6) and (7).

$$V_1 = (\frac{|a_{11}|}{|R_1|}, \frac{|a_{12}|}{|R_1|}, \cdots, \frac{|a_{mk}|}{|R_m|}) \tag{6}$$

where $m$ indicates the number of groups whose value is 4 according to DHP. $|R_i|$ indicates the frequency of $R_i$. $|a_{mk}|$ indicates the frequency of $k$-$th$ character in the $m$-$th$ kind of group of the reduced sequence. The dimension of the vector $V_1$ is $20 \times 6 = 120$.

The encoding process of the vector $V_2$ is shown as

$$V_2 = (\frac{|R_1R_1|}{|R_1|}, \frac{|R_1R_2|}{|R_1|}, \cdots, \frac{|R_mR_j|}{|R_m|}) \tag{7}$$

where $|\mathrm{R}_m R_j|$ indicates the frequency of di-character $R_m R_j$. The dimension of the feature vector $V_2$ is $5^2 + 3^3 + 3^2 + 3^2 + 3^2 + 4^2 = 77$. We can obtain the total dimension $120+77=197$ by fusing vector $V_1$ and $V_2$.

*Autocorrelation descriptors*

In GTB-PPI model, three autocorrelation descriptors (AD) are selected: Morean-Broto autocorrelation (NMBA), Moran autocorrelation (MA), Geary autocorrelation (GA) [6]. The values of the seven physicochemical properties corresponding to the 20 amino acids are shown in Table S3.

$$NMBA(l) = \frac{MBA(l)}{N - l}, \ l = 1, 2, \cdots, lag \tag{8}$$

where $MBA(l) = \sum_{i=1}^{N-l} P(AA_i)P(AA_{i+l})$. $P(AA_i)$ and $P(AA_{i+l})$ are standardized property values at the position of amino acid $AA_i$ and $AA_{i+l}$, respectively. The $lag$ is the parameter needs to be adjusted.

$$MA(l) = \frac{\frac{1}{N-l}\sum_{i=1}^{N-l}(P(AA_i) - \mu)(P(AA_{i+l}) - \mu)}{\frac{1}{N}\sum_{i=1}^{N}(P(AA_i) - \mu)^2}, \ l = 1, 2, \cdots, lag \tag{9}$$

where $\mu$ is the mean value of each amino acid.

$$GA(l) = \frac{\frac{1}{2(N-l)}\sum_{i=1}^{N-l}(P(AA_i) - P(AA_{i+l}))^2}{\frac{1}{N}\sum_{i=1}^{N}(P(AA_i) - \mu)^2}, l = 1, 2, \cdots, lag \quad (10)$$

We can obtain $3\times 7\times lag$ features from protein sequence using autocorrelation descriptor, where $lag$ is the interval of the amino acid residues, 3 represents three autocorrelation descriptors, 7 represents seven physicochemical properties.


#### File S2 The parameter selection of $\xi$ and $lag$

For the selection of parameter $\xi$ in PsePSSM, $\xi$ values are different when two datasets achieve the highest overall prediction accuracy. The peak point of the *H.*

*pylori* dataset is 3, and the optimal value $\xi$ in the *S. cerevisiae* dataset is 9. We can obtain important evolutional information from PsePSSM, and the *S. cerevisiae* dataset is selected as train dataset to test independent dataset. Thus, the parameter $\xi$ is set as nine in PsePSSM to extract the evolutionary information. The parameters in AD that reached the peak value are the same, all of which are 11. Based on above discussion, the PseAAC in the GTB-PPI model is set to 11, the PsePSSM is set to 9, and the AD is set to 11 in the model.

**File S3 The parameter optimization of L1-RLR**

Although feature fusion can acquire important, valuable feature information in the process of PPIs prediction, the increase of dimension inevitably generates some unimportant features for classification. This case could have adverse effects on PPIs prediction. According to Equation (1), different penalty parameter $C$ values can be selected to determine the different subset. The penalty $C$ is set as 0.1, 0.3, 0.5, 0.8, 1, 1.2, 1.5, and 2 respectively and the GTB classifier is employed to predict PPIs via five-fold cross-validation (Table S7). The number of original features and selected feature subsets of PseAAC, PsePSSM, RSIV, and AD are shown in Figure S1 and Figure S2. From Table S7, on *S. cerevisiae* and *H. pylori*, GTB-PPI achieve the best prediction performance $C$=1. When the $C$ value is 1, 331 optimal features are selected for *S. cerevisiae,* and 199 optimal features have been remained for *H. pylori*.

**File S4 The comparison of dimensional-reduction methods**

From Figure 3C, in PR curves, L1-RLR almost obtains the higher precision value at corresponding recall value. For the *S. cerevisiae*, the AUPR values of L1-RLR, SSDR, PCA, KPCA, FA, mRMR, and CMIM are 0.9847, 0.9388, 0.9386, 0.9271, 0.9510, 0.9728, and 0.9717, respectively. The AUPR value of L1-RLR is 1.19%-5.76% higher than the other six dimensional-reduction methods. From Figure 3D, for *H. pylori*, the L1-RLR achieves the best performance. The AUPR values of L1-RLR, SSDR, PCA, KPCA, FA, mRMR, and CMIM are 0.9498, 0.9077, 0.8485, 0.8728, 0.8665, 0.9464, and 0.8516, respectively. The AUPR value of L1-RLR is 0.34%-10.13% higher than six dimensional-reduction methods. Thus, the results indicate L1-RLR can effectively reduce redundant information and achieve excellent performance in predicting PPIs.

**File S5 The comparison of different classifiers**

From Figure 4A, the ROC curves show that the GTB classifier outperforms the prediction results of other four classifier methods. For *S. cerevisiae* dataset, the AUC values for KNN, NB, SVM, RF and GTB are 0.9172, 0.7922, 0.9618, 0.9762, 0.9875, respectively. The AUC value of GTB is 7.03% higher than KNN and 2.57% higher than SVM. From Figure 4B, we can see GTB is superior to other classifiers for *H. pylori* dataset. The AUC values for KNN, NB, SVM, RF and GTB are 0.8683, 0.7775, 0.9214, 0.9509, 0.9559, respectively. The GTB is 3.45% higher than SVM for AUC value. From Figure 4C, the PR curves show that GTB is superior to the other classifiers for PPIs prediction. Compared with other dimensional reduction methods, L1-RLR achieved the better precision almost at every recall value. For *S. cerevisiae* dataset, the AUPR values of KNN, NB, SVM, RF, and GTB are 0.9006, 0.7921, 0.9581, 0.9709 and 0.9847, respectively. The AUPR value of GTB is 8.41%, 19.26%, 2.66%, and 1.38% higher than KNN, NB, SVM and RF respectively. Figure 4D plots the prediction performance of different classifiers on the *H. pylori* dataset, and the AUPR values of KNN, NB, SVM, RF, and GTB are 0.8531, 0.7625, 0.9182, 0.9477, 0.9498, respectively. The AUPR value of GTB is 3.16% higher than that of SVM. The GTB classifier has the best prediction performance for PPIs and has better generalization performance by analyzing the two datasets.

### Supplementary figures

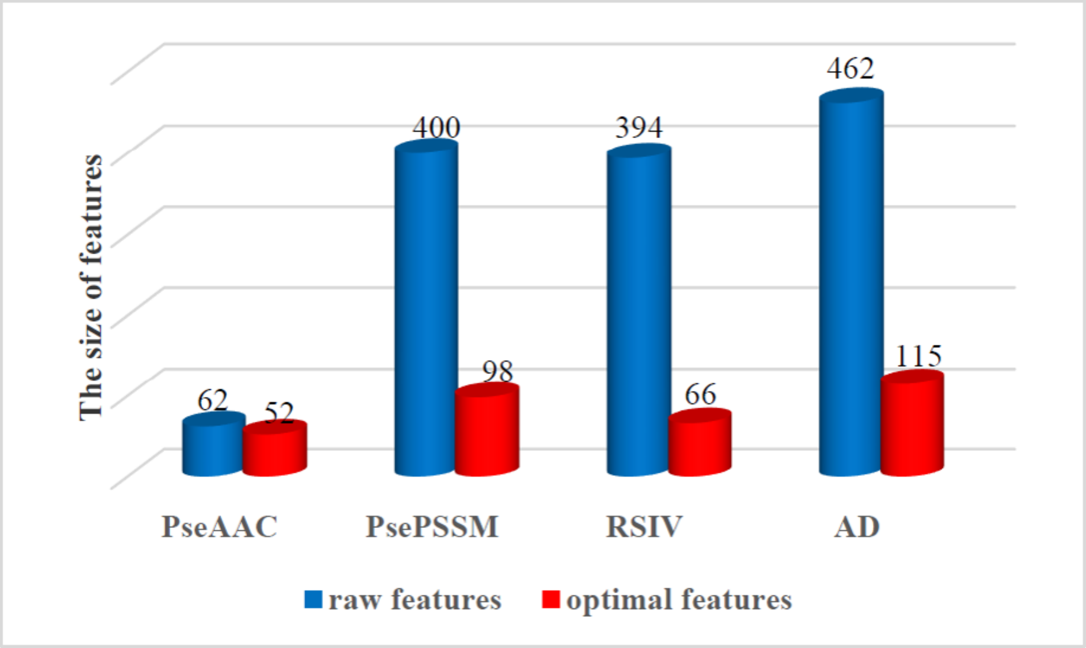


**Figure S1 The number of feature subset with different feature extraction methods on *S. cerevisiae* dataset**

The raw features represent the number of initial feature dimension and the optimal features represent the number of the selected features.

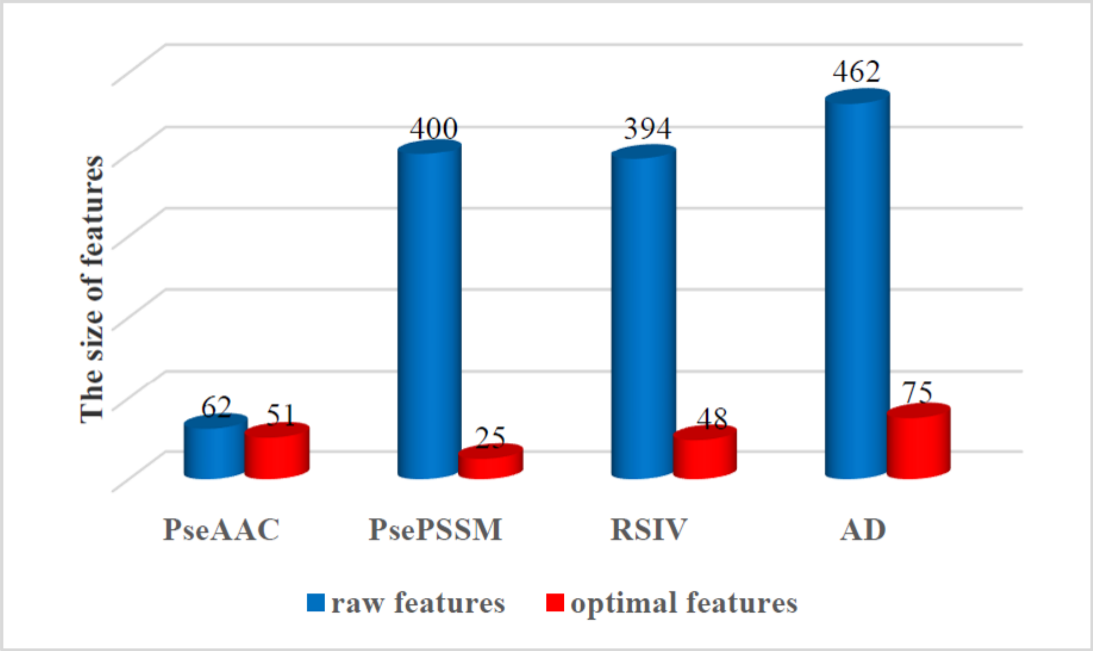


**Figure S2 The number of feature subset with different feature extraction**

**methods on *H. pylori* dataset**

The raw features represent the number of initial feature dimension and the optimal features represent the number of the selected features.

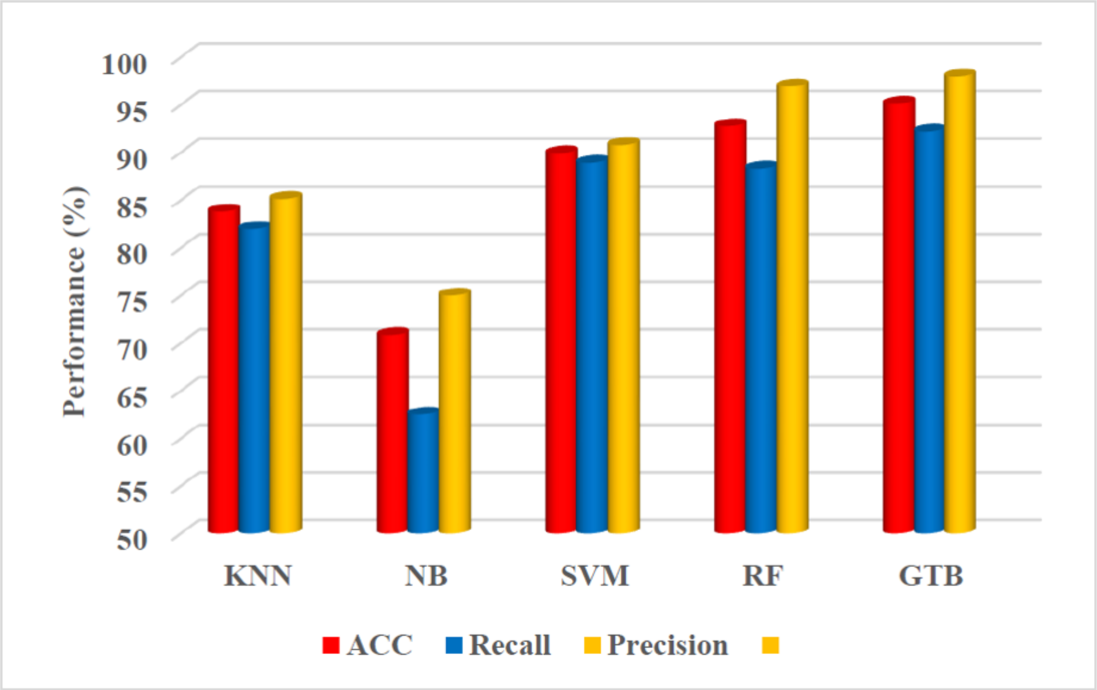


**Figure S3 The comparison of prediction results with different classifiers on *S. cerevisiae* dataset**

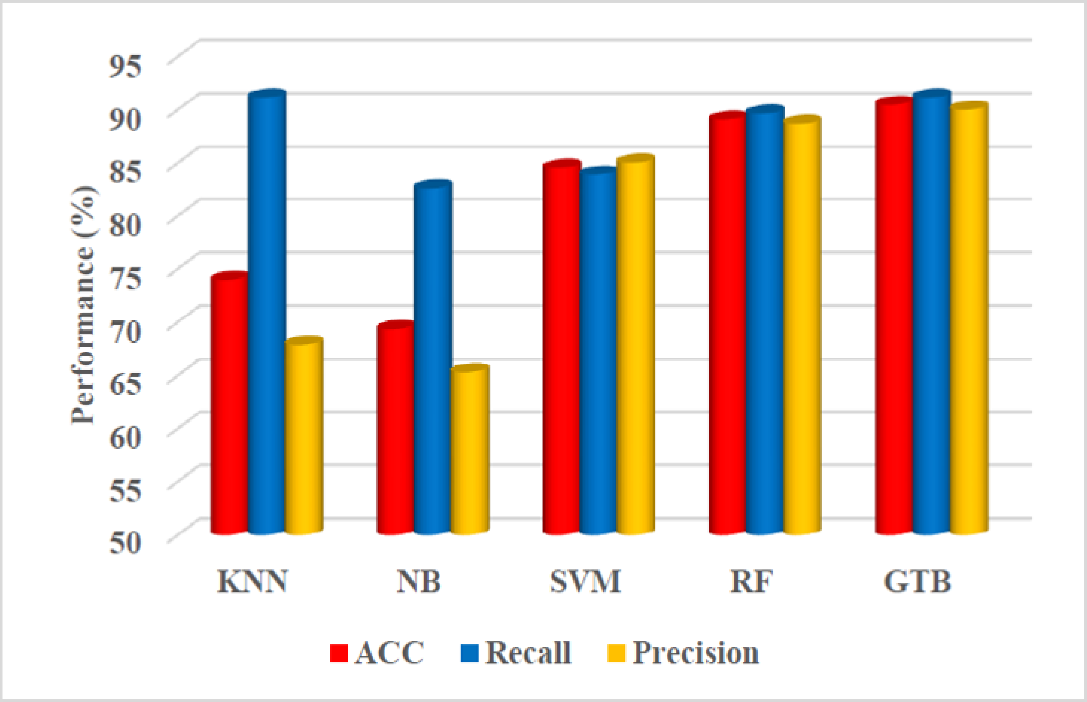

**Figure S4 The comparison of prediction results with different classifiers on *H. pylori* dataset**

### Supplementary tables

**Table S1 The unique proteins for each dataset**

| Dataset | Protein pairs | Unique proteins |
|---|---|---|
| *S. cerevisiae* | 11,188 | 2530 |
| *H. pylori* | 2,916 | 1428 |
| *C. elegans* | 4,013 | 2629 |
| *E. coli* | 6,954 | 1830 |
| *H. sapiens* | 1,412 | 1083 |
| *M. musculus* | 313 | 355 |

**Table S2 Amino acid classification according to different physicochemical property**

| Property | Classifications |
|---|---|
| Polarity/acidity | DE RHK WYF SCMNQT GAVLIP |
| Acidity | DE KHR ACFGILMNPQSTVWY |
| Secondary structure | EHALMQKR VTIYCWF GDNPS |
| Charge | KR AVNCQGHILMFPSTWY DE |
| DHP | PALVIFWM QSTYCNG HKR DE |
| Hydrophobicity | RKEDQN GASTPHY CLVIMFW |

**Table S3 The original values of the seven physicochemical properties for the 20 native amino acids**

| Amino acid | $\phi^{(1)}$ | $\phi^{(2)}$ | $\phi^{(3)}$ | $\phi^{(4)}$ | $\phi^{(5)}$ | $\phi^{(6)}$ | $\phi^{(7)}$ |
|---|---|---|---|---|---|---|---|
| A | 0.62 | -0.5 | 27.5 | 8.1 | 0.046 | 1.181 | 0.007187 |
| C | 0.29 | -1 | 44.6 | 5.5 | 0.128 | 1.461 | -0.03661 |
| D | -0.9 | 3 | 40 | 13 | 0.105 | 1.587 | -0.02382 |
| E | -0.74 | 3 | 62 | 12.3 | 0.151 | 1.862 | 0.006802 |
| F | 1.19 | -2.5 | 115.5 | 5.2 | 0.29 | 2.228 | 0.03755 |
| G | 0.48 | 0 | 0 | 9 | 0 | 0.881 | 0.1791 |
| H | -0.4 | -0.5 | 79 | 10.4 | 0.23 | 2.025 | -0.01069 |
| I | 1.38 | -1.8 | 93.5 | 5.2 | 0.186 | 1.81 | 0.02163 |
| K | -1.5 | 3 | 100 | 11.3 | 0.219 | 2.258 | 0.01771 |
| L | 1.06 | -1.8 | 93.5 | 4.9 | 0.186 | 1.931 | 0.05167 |
| M | 0.64 | -1.3 | 94.1 | 5.7 | 0.221 | 2.034 | 0.002683 |
| N | -0.78 | 2 | 58.7 | 11.6 | 0.134 | 1.655 | 0.005392 |
| P | 0.12 | 0 | 41.9 | 8 | 0.131 | 1.468 | 0.2395 |
| Q | -0.85 | 0.2 | 80.7 | 10.5 | 0.18 | 1.932 | 0.04921 |
| R | -2.53 | 3 | 105 | 10.5 | 0.291 | 2.56 | 0.04359 |
| S | -0.18 | 0.3 | 29.3 | 9.2 | 0.062 | 1.298 | 0.004627 |
| T | -0.05 | -0.4 | 51.3 | 8.6 | 0.108 | 1.525 | 0.003352 |
| V | 1.08 | -1.5 | 71.5 | 5.9 | 0.14 | 1.645 | 0.057 |

| | | | | | | | |
|---|---|---|---|---|---|---|---|
| W | 0.81 | -3.4 | 145.5 | 5.4 | 0.409 | 2.663 | 0.03798 |
| Y | 0.26 | -2.3 | 117.3 | 6.2 | 0.298 | 2.368 | 0.0236 |

*Note*: $\phi(1)$ is hydrophobicity; $\phi(2)$ is hydrophilicity; $\phi(3)$ is side chain volume; $\phi(4)$ is polarity; $\phi(5)$ is polarizability; $\phi(6)$ is solvent-accessible surface area and $\phi(7)$ is side chain net charge index.

**Table S4 Performance comparison with different $\lambda$ values on PPIs datasets**

| Dataset | Evaluation | $\lambda$ | | | | | |
|---|---|---|---|---|---|---|---|
| | | **1** | **3** | **5** | **7** | **9** | **11** |
| *S. cerevisiae* | ACC | 94.38 | 94.57 | 94.59 | 94.50 | **94.87** | 94.60 |
| | Recall | 91.99 | 91.96 | 92.06 | 92.08 | 92.42 | 92.08 |
| | Precision | 96.61 | 97.02 | 96.97 | 96.78 | 97.18 | 96.98 |
| | MCC | 0.8886 | 0.8926 | 0.8930 | 0.8911 | 0.8985 | 0.8933 |
| *H. pylori* | ACC | 88.72 | 88.72 | 88.41 | 89.09 | 89.16 | **89.44** |
| | Recall | 87.72 | 87.45 | 86.83 | 87.92 | 88.75 | 88.89 |
| | Precision | 89.60 | 89.78 | 89.67 | 90.08 | 89.49 | 89.90 |
| | MCC | 0.7753 | 0.7751 | 0.7686 | 0.7827 | 0.7833 | 0.7890 |

**Table S5 Performance comparison with different $\xi$ values on PPIs datasets**

| Dataset | Evaluation | $\xi$ | | | | | |
|---|---|---|---|---|---|---|---|
| | | **1** | **3** | **5** | **7** | **9** | **11** |
| *S. cerevisiae* | ACC | 94.58 | 94.71 | 94.87 | 94.68 | **94.98** | 94.75 |
| | Recall | 92.08 | 92.08 | 92.31 | 92.06 | 92.47 | 92.08 |
| | Precision | 96.94 | 97.19 | 97.29 | 97.15 | 97.37 | 97.28 |
| | MCC | 0.8928 | 0.8955 | 0.8986 | 0.8949 | 0.9009 | 0.8964 |
| *H. pylori* | ACC | 87.18 | **88.85** | 88.62 | 88.00 | 88.20 | 88.75 |
| | Recall | 86.97 | 89.02 | 88.96 | 88.00 | 88.96 | 89.09 |
| | Precision | 87.34 | 88.74 | 88.43 | 88.05 | 87.74 | 88.49 |
| | MCC | 0.7436 | 0.7773 | 0.7729 | 0.7603 | 0.7648 | 0.7751 |

**Table S6 Prediction results with the different $lag$ value on PPIs data set**

| Dataset | Evaluation | $lag$ | | | | | |
|---|---|---|---|---|---|---|---|
| | | **1** | **3** | **5** | **7** | **9** | **11** |
| *S. cerevisiae* | ACC | 93.73 | 94.12 | 94.52 | 94.37 | 94.32 | **94.53** |
| | Recall | 91.37 | 91.20 | 91.72 | 91.33 | 91.29 | 91.47 |
| | Precision | 95.91 | 96.85 | 97.16 | 97.25 | 97.18 | 97.43 |
| | MCC | 0.8757 | 0.8839 | 0.8918 | 0.8890 | 0.8880 | 0.8923 |

| | | | | | | | |
|---|---|---|---|---|---|---|---|
| *H. pylori* | ACC | 85.60 | 88.30 | 89.03 | 89.44 | 89.10 | **89.68** |
| | Recall | 84.43 | 86.70 | 88.20 | 88.75 | 88.13 | 89.44 |
| | Precision | 86.50 | 89.58 | 89.68 | 90.01 | 89.95 | 89.92 |
| | MCC | 0.7126 | 0.7667 | 0.7809 | 0.7890 | 0.7828 | 0.7938 |

**Table S7 Effect of selecting different penalty parameter on the model performance**

| Dataset | Evaluation | *C* | | | | | | | |
|---|---|---|---|---|---|---|---|---|---|
| | | **0.1** | **0.3** | **0.5** | **0.8** | **1** | **1.2** | **1.5** | **2** |
| *S. Cerevisiae* | ACC | 94.45 | 94.94 | 94.91 | 95.10 | **95.15** | 94.97 | 94.88 | 95.03 |
| | Recall | 91.58 | 92.06 | 92.24 | 92.24 | 92.21 | 92.21 | 92.03 | 92.22 |
| | Precision | 97.16 | 97.69 | 97.43 | 97.43 | 97.97 | 97.60 | 97.59 | 97.71 |
| | MCC | 0.8905 | 0.9003 | 0.8944 | 0.8944 | 0.9045 | 0.9008 | 0.8991 | 0.9026 |
| *H. pylori* | ACC | 89.16 | 89.75 | 89.81 | 89.61 | **90.47** | 89.95 | 89.95 | 89.82 |
| | Recall | 88.96 | 89.98 | 89.30 | 88.96 | 89.99 | 89.85 | 89.78 | 89.78 |
| | Precision | 89.32 | 89.60 | 90.23 | 90.17 | 91.15 | 90.10 | 90.09 | 89.88 |
| | MCC | 0.7833 | 0.7954 | 0.7961 | 0.7925 | 0.8100 | 0.7996 | 0.7991 | 0.7966 |

**Table S8 Performance of kernel principle component analysis with different contribution rate**

| Dataset | Evaluation | The rate of contribution (%) | | | |
|---|---|---|---|---|---|
| | | **80** | **85** | **90** | **95** |
| *S. cerevisiae* | ACC | 85.49 | 85.37 | 85.36 | **85.63** |
| | Recall | 84.93 | 85.15 | 84.97 | 85.23 |
| | Precision | 85.90 | 85.52 | 85.65 | 85.94 |
| | MCC | 0.7100 | 0.7075 | 0.7073 | 0.7127 |
| *H. pylori* | ACC | 79.08 | 78.88 | **80.42** | 78.50 |
| | Recall | 77.16 | 78.19 | 79.15 | 77.92 |
| | Precision | 80.26 | 79.27 | 81.31 | 78.91 |
| | MCC | 0.5823 | 0.5776 | 0.6092 | 0.5710 |

**Table S9 Performance of principle component analysis with different contribution rate**

| Dataset | Evaluation | The rate of contribution (%) | | | |
|---|---|---|---|---|---|
| | | **80** | **85** | **90** | **95** |
| *S. cerevisiae* | ACC | 86.16 | 86.59 | 86.91 | **87.62** |
| | Recall | 85.63 | 85.88 | 86.04 | 86.29 |
| | Precision | 86.56 | 87.12 | 87.58 | 88.65 |
| | MCC | 0.7234 | 0.7320 | 0.7384 | 0.7527 |

| | | | | | |
|---|---|---|---|---|---|
| *H. pylori* | ACC | 79.42 | 79.80 | **80.18** | 79.25 |
| | Recall | 77.09 | 78.67 | 77.91 | 76.48 |
| | Precision | 80.90 | 80.53 | 81.60 | 80.98 |
| | MCC | 0.5893 | 0.5964 | 0.6043 | 0.5861 |

**Table S10 Comparison of prediction results on different dimensional reduction methods**

| **Dataset** | **Evaluation** | **Method** | | | | | | |
|---|---|---|---|---|---|---|---|---|
| | | **L1-RLR** | **SSDR** | **PCA** | **KPCA** | **FA** | **mRMR** | **CMIM** |
| *S. cerevisiae* | ACC | 95.15 | 87.02 | 87.62 | 85.63 | 88.94 | 92.55 | 92.32 |
| | Recall | 92.21 | 85.47 | 86.29 | 85.23 | 87.20 | 90.01 | 89.65 |
| | Precision | 97.97 | 88.26 | 88.65 | 85.94 | 90.36 | 94.84 | 94.74 |
| | MCC | 0.9045 | 0.7410 | 0.7527 | 0.7127 | 0.7794 | 0.8523 | 0.8479 |
| *H. pylori* | ACC | 90.47 | 85.25 | 79.25 | 78.50 | 80.28 | 87.76 | 78.43 |
| | Recall | 89.99 | 77.16 | 76.47 | 77.92 | 77.02 | 87.59 | 78.33 |
| | Precision | 91.15 | 92.12 | 80.98 | 78.91 | 82.40 | 87.91 | 78.54 |
| | MCC | 0.8100 | 0.7151 | 0.5861 | 0.5710 | 0.6073 | 0.7553 | 0.5691 |

**Table S11 The AUC and AUPR on different dimensional reduction methods**

| **Dataset** | **Evaluation** | **L1-RLR** | **SSDR** | **PCA** | **KPCA** | **FA** | **mRMR** | **CMIM** |
|---|---|---|---|---|---|---|---|---|
| *S. cerevisiae* | AUC | 0.9875 | 0.9445 | 0.9420 | 0.9305 | 0.9568 | 0.9770 | 0.9769 |
| | AUPR | 0.9847 | 0.9388 | 0.9386 | 0.9271 | 0.9510 | 0.9728 | 0.9717 |
| *H. pylori* | AUC | 0.9559 | 0.9238 | 0.8706 | 0.8803 | 0.8824 | 0.9461 | 0.8726 |
| | AUPR | 0.9498 | 0.9077 | 0.8485 | 0.8728 | 0.8665 | 0.9464 | 0.8516 |

**Table S12 Performance of K nearest neighbor with different size of neighbor**

| Dataset | Evaluation | The size of neighbor | | | | | | |
|---|---|---|---|---|---|---|---|---|
| | | 1 | 3 | 5 | 7 | 9 | 15 | 20 |
| *S. cerevisiae* | ACC | 82.40 | 83.74 | **83.82** | 83.39 | 83.17 | 81.89 | 81.43 |
| | Recall | 81.09 | 81.69 | 81.98 | 82.17 | 82.77 | 83.57 | 83.68 |
| | Precision | 83.29 | 85.18 | 85.13 | 84.24 | 83.46 | 80.86 | 80.10 |
| | MCC | 0.6484 | 0.6754 | 0.6770 | 0.6682 | 0.6638 | 0.6384 | 0.6295 |
| *H. pylori* | ACC | **74.63** | 73.77 | 73.97 | 72.43 | 71.33 | 71.06 | 70.96 |
| | Recall | 88.55 | 91.02 | 91.15 | 90.95 | 90.60 | 91.02 | 91.15 |
| | Precision | 69.28 | 67.69 | 67.86 | 66.41 | 65.42 | 65.16 | 64.99 |
| | MCC | 0.5129 | 0.5064 | 0.5106 | 0.4831 | 0.4625 | 0.4594 | 0.4580 |

**Table S13 Performance of random forest with different size of base decision tree**

| Dataset | Evaluation | The size of base decision tree | | | | |
|---|---|---|---|---|---|---|
| | | 50 | 100 | 500 | 1000 | 2000 |
| *S. cerevisiae* | ACC | 92.69 | 92.70 | 92.78 | 92.79 | **92.84** |
| | Recall | 88.24 | 88.38 | 88.34 | 88.29 | 88.38 |
| | Precision | 96.86 | 96.74 | 96.94 | 97.04 | 97.04 |
| | MCC | 0.8572 | 0.8572 | 0.8590 | 0.8594 | 0.8603 |
| *H. pylori* | ACC | 87.76 | 88.72 | **89.06** | 88.75 | 88.72 |
| | Recall | 88.41 | 89.78 | 89.64 | 89.71 | 89.44 |
| | Precision | 87.30 | 87.96 | 88.67 | 88.06 | 88.21 |
| | MCC | 0.7554 | 0.7749 | 0.7818 | 0.7754 | 0.7748 |

**Table S14 Prediction results of different classifiers on S. *cerevisiae*, *H. pylori* dataset**

| Dataset | Model | ACC (%) | Recall (%) | Precision (%) | MCC |
|---|---|---|---|---|---|
| *S. cerevisiae* | KNN | 83.82±0.64 | 81.98±0.80 | 85.13±1.04 | 0.6770±0.0131 |
| | NB | 70.86±1.81 | 62.53±3.18 | 75.01±1.60 | 0.4233±0.0351 |
| | SVM | 89.94±0.44 | 88.93±0.65 | 90.77±0.59 | 0.7991±0.0088 |
| | RF | 92.78±0.48 | 88.34±0.61 | 96.94±0.64 | 0.8590±0.0096 |
| | GTB | 95.15±0.25 | 92.21±0.36 | 97.97±0.60 | 0.9045±0.0053 |
| *H. pylori* | KNN | 73.97±1.87 | 91.15±1.65 | 67.86±1.61 | 0.5106±0.0372 |
| | NB | 69.34±1.42 | 82.58±0.79 | 65.32±1.47 | 0.4012±0.0267 |
| | SVM | 84.54±1.53 | 83.88±2.08 | 85.05±2.22 | 0.6912±0.0311 |

| | | | | | |
|---|---|---|---|---|---|
| | RF | 89.06±0.81 | 89.64±1.77 | 88.67±1.72 | 0.7818±0.0161 |
| | GTB | 90.47±0.84 | 91.15±1.42 | 89.99±2.06 | 0.8100±0.0163 |

**Table S15 The AUC and AUPR of different classifiers on S. *cerevisiae*, *H. pylori* dataset**

| **Dataset** | **Evaluation** | **Method** | | | | |
|---|---|---|---|---|---|---|
| | | **KNN** | **NB** | **SVM** | **RF** | **GTB** |
| *S. cerevisiae* | AUC | 0.9172 | 0.7922 | 0.9618 | 0.9762 | 0.9875 |
| | AUPR | 0.9006 | 0.7921 | 0.9581 | 0.9709 | 0.9847 |
| *H. pylori* | AUC | 0.8683 | 0.7775 | 0.9214 | 0.9509 | 0.9559 |
| | AUPR | 0.8531 | 0.7625 | 0.9182 | 0.9477 | 0.9498 |
